# Continuous motor skills as flexible control policies: a video game study

**DOI:** 10.1101/2023.09.21.558913

**Authors:** David M. Huberdeau, Adrian M. Haith, John W. Krakauer

## Abstract

Many motor skills consist of continuous sequential actions, such as a tennis serve. It is currently unclear how these surprisingly understudied behaviors are learned, with the leading hypothesis being that sequences of single actions become “chunked” into larger single executable units. Under this hypothesis, continuous sequential actions should become more task-specific and less generalizable with practice. To test this, we developed a video game that requires participants to hold a tablet with both hands and steer a virtual car (the “ant car”) along a curving track. We tested participants’ ability to generalize their skill to a probe track that required a different sequence of turns. Across days of practice, task success increased, and movement variability decreased. On the probe track, movement quality at the level of kinematics fully generalized but performance at the level of task success showed a consistent decrement. To address this apparent paradox, we empirically derived the control policy participants used at their maximal skill level on the training track. Notably, this policy was fully transferred to the probe track, but there were more instances of momentary deviations from it (lapses), which explains the worse performance despite equivalent skill. We conclude that continuous motor skills are acquired through learning of a flexible control policy that maps states onto actions and not through chunking or automatizing of a specific sequence of actions.

## Introduction

Many human motor skills require precise execution of continuous sequential actions. For example, a tennis serve is made up of subcomponent movements that blend seamlessly, and each subcomponent is also made up of a continuous sequence of joint rotations and muscle activations. Learning to better execute continuous sequential actions is less well studied than learning to produce discrete sequences of actions (Verwey, 2001; Diedrichsen and Kornysheva, 2015; Hardwick et al., 2019). Discrete sequence learning paradigms emphasize learning to select the right actions rather than improving movement execution of each individual action, which are usually over-learned. For example, in a sequence of button presses, each button press is itself easy.

One theory of how performance improves as discrete sequences are learned is that individual sequence elements are grouped into longer sequence fragments (often called chunks), which can then be executed as a single unit (Povel and Collard, 1982; Berns and Sejnowski, 1998; David A. Rosenbaum et al., 2001; Yamaguchi and Logan, 2014; Diedrichsen and Kornysheva, 2015). Recent work suggests that chunking in discrete sequence tasks is primarily related to cognition rather than to motor execution, which is to say that it is knowledge of the order of actions that is chunked rather than the commands for their execution (Diedrichsen and Kornysheva, 2015; Wong et al., 2015; Zimnik and Churchland, 2021). For example, knowing one’s ATM number as a single unit may exist at a cognitive level, with this order then communicated to motor cortex one element at a time for execution. In this example, no motor chunking is necessary.

Critically, what distinguishes continuous sequential actions from discrete sequences of actions is that improvement occurs at the level of movement execution – a tennis player’s forehand gets faster and more accurate. Chunking at the knowledge level for such continuous sequential actions could be a challenge, as there are no clear divides between movement elements. It is therefore interesting to consider the possibility that chunking does occur for these tasks, but at the level of movement execution rather than action selection.

In a previous study to address how continuous sequential actions improve at the level of movement execution, we developed a wrist-controlled cursor task (the arc-pointing task) that required participants to control a cursor to make a fast trajectory through a U-shaped channel without touching or crossing its edges (Shmuelof et al., 2012). Participants learned to get through the channel, a binary outcome, with increasing success. Additionally, continuous trajectory kinematics became smoother and less variable, and feedback control improved, leading to improved motor execution and greater task success. Other studies of continuous sequential actions have also shown shifts in the speed-accuracy trade-off function (Reis et al., 2009). The improvements in movement execution seen in the arc-pointing task, what we called “motor acuity”, could arguably have been due either to chunking of a sequence of actions or by learning a de novo control policy that maps a sequence of states onto actions (Telgen et al., 2014; Yang et al., 2021; Hadjiosif et al., 2022).

To distinguish whether continuous sequential actions are learned by means of a continuous control policy or by chunking a sequence of actions, and to better understand how the quality of execution improves with practice, we created a novel tablet computer-based video game (Figure 1a) that requires continuous bilateral movements of the arms and wrists to steer a virtual car along a narrow, curved track (Figure 1b) at constant speed. Participants practiced the game for 1-, 3-, 5-, or 10-days prior to a “probe” of generalization (Figure 1d) on the mirrored version of the track (figure 1c). Successful generalization of this skill to the novel kinematics required on the mirrored track would be inconsistent with having learned to chunk a fixed sequence of actions (see Supplemental Figure 1).

**Figure 1:**
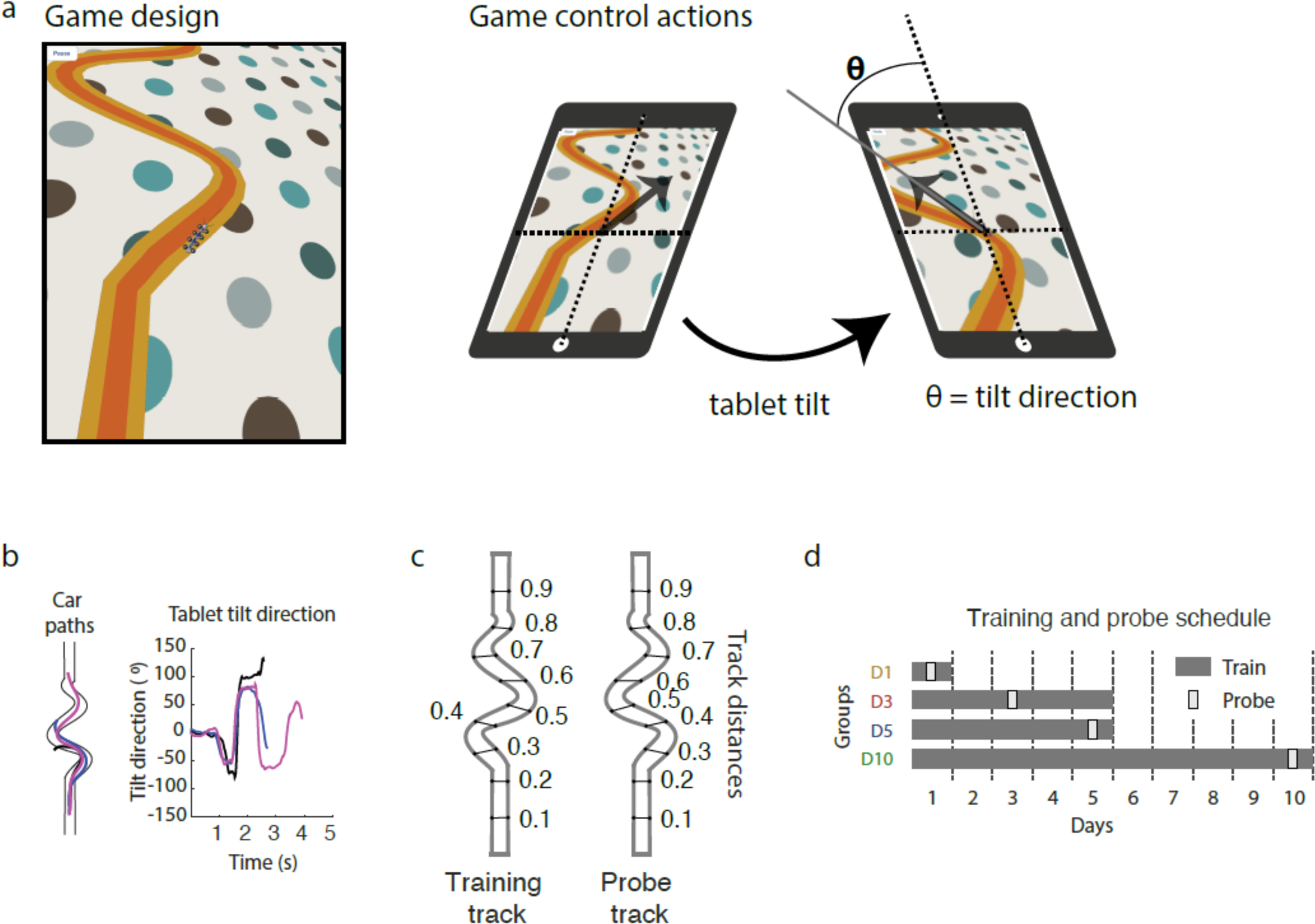
The ant-car game.. a, The game’s artistic design and control actions. The car was styled like an insect and the track was demarcated by an orange ribbon on a polka-dotted background. The roll and pitch of the tablet computer determined the direction of acceleration of the car. b, Sample recordings of the trajectory of the car and the direction of the tablet tilt. c, Training and Probe tracks. d, Training and Probe trial assignments per group. Groups trained for varying numbers of days (grey bars), up to a maximum of ten days, and were probed for generalization at different times throughout learning (white boxes).

Introducing the probe on different days of training across the four groups allowed us to ascertain whether generalization properties changed as the attained skill level increased.

## Results

Participants practiced a novel video game for up to two weeks that required navigating a cartoon car (the “ant-car”) along a narrow and winding track (see methods for details). After practice, participants became able to travel further along the track without falling off, generating smoother and more consistent ant-car (Figure 2a) and tablet tilt (Figure 2b) trajectories. The distance travelled along the track increased with practice. Linear models fit to performance in windows of 25 trials (Figure 3a) demonstrated significant changes in average distance travelled across practice for each group (Linear regression; D1: F(82) = 25.38, p < .001; D3: F(605) = 288.4, p < 0.001; D5: F(757) = 122.1, p < 0.001; D10: F(1474) = 402.4, p < 0.001).

**Figure 2:**
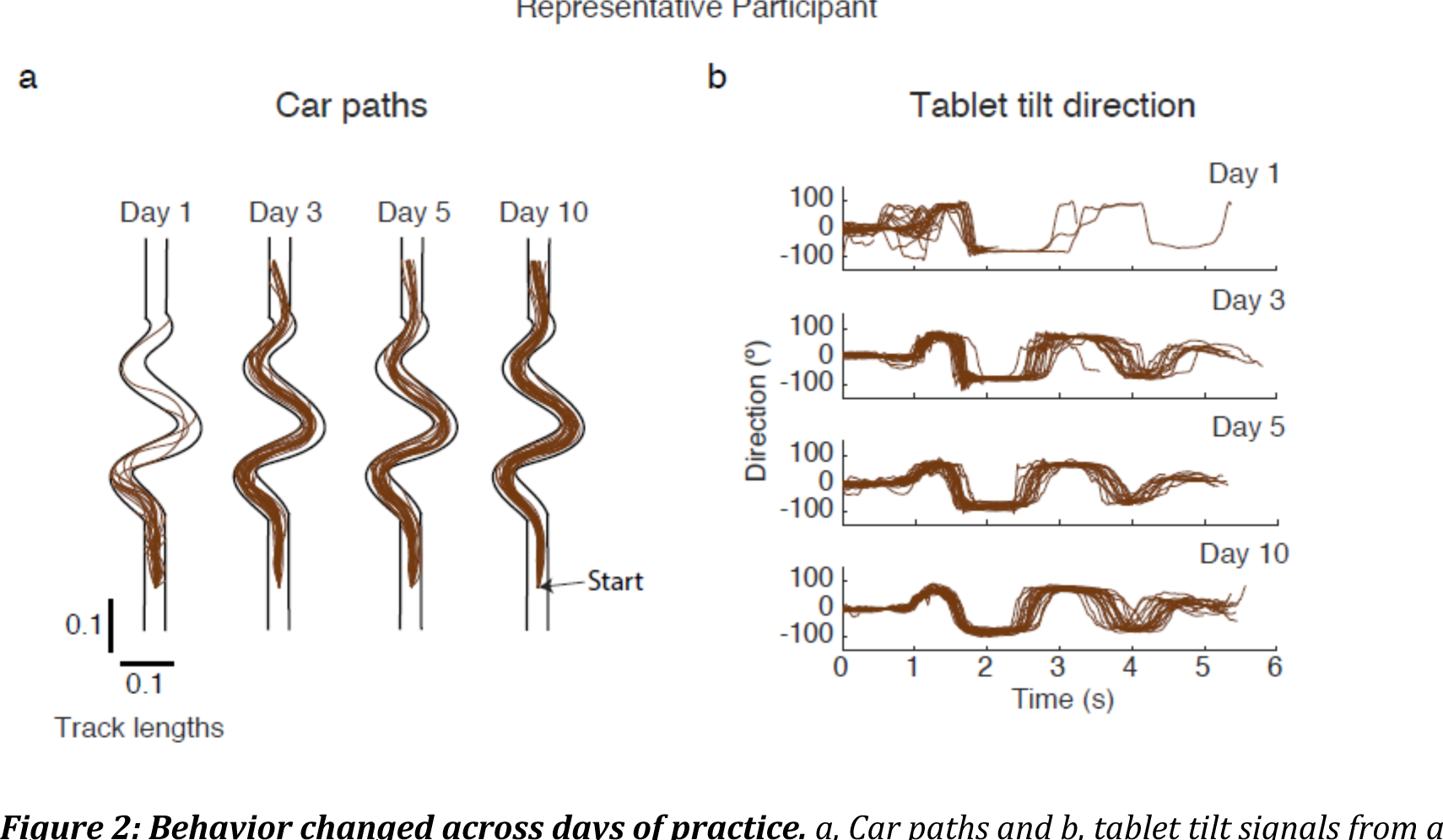
Behavior changed across days of practice. a, Car paths and b, tablet tilt signals from a representative participant from group D10.

**Figure 3:**
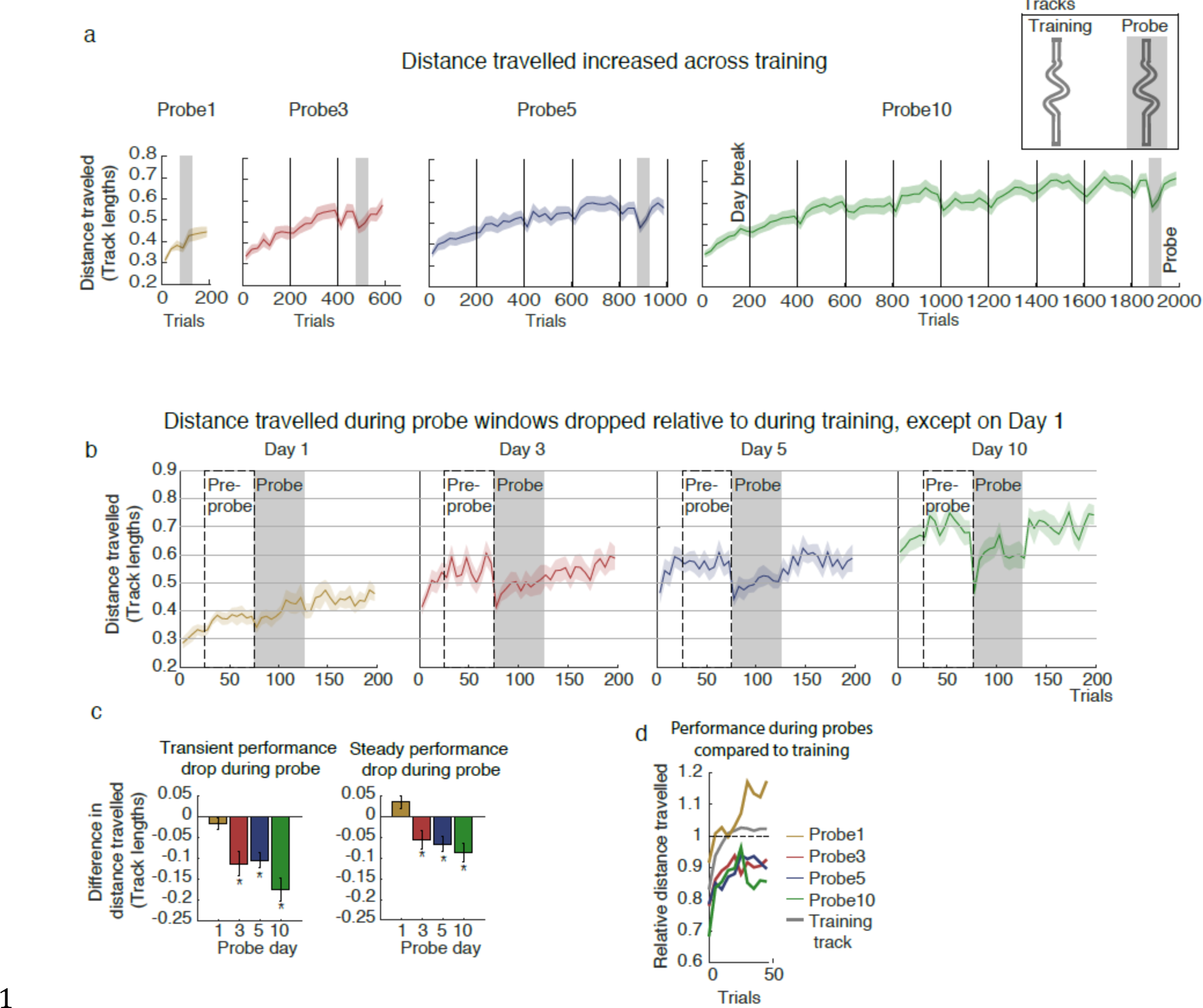
The distance traveled on the Training track increased with practice but decreased in a consistent way during Probes after Day 1. a, The mean distance travelled along the path within bins of 25 trials, averaged across participants (mean ± standard error) for each group. Vertical lines indicate overnight breaks. Grey bars indicate a block of Probe trials. b, The distance travelled in bins of five trials and averaged across participants (mean ± standard error) during days when a Probe was encountered. c, The average mean ± standard error of the distance travelled in the first five trials after the onset of a block of Probe trials (left panel) and in the final 45 trials of a block of Probe trials (right panel). d, The average distance travelled in blocks of five trials relative to the previous 50 trials for a day of training trials (grey trace) and for each block of probe trials (yellow, red, green, and blue traces).

At different points during learning, each group performed a series of probe trials in which the layout of the track was mirror-reversed compared to training. For participants who experienced the probe during the first day of training (Group D1), performance during the probe trials was comparable to the immediately preceding training trials. In contrast, for participants probed during days 3, 5, or 10 of training (Groups D3, D5, and D10), the distance travelled decreased during probe trials compared to the pre-probe window (Figure 3b). This performance decrease in the probe trials had two components: an immediate drop during the initial five trials (ANOVA: F(3) = 7.06, p < 0.001; t-tests: D1: t = 1.34, p = 0.19; D3: t = -3.05, p < 0.01; D5: t = -3.55, p < 0.01; D10: t = -4.44, p < 0.001) followed by recovery to an asymptote that remained approximately constant for the remainder of the probe window (Figure 3c & d).

For the asymptotic period, defined as the final 45 trials of the probe block, the difference in distance travelled between the probe and pre-probe windows was significantly different among groups (ANOVA: F(3) = 9.59, p < 0.001), and groups D3, D5, and D10 had significantly lower distance travelled during probes (t-tests: D1: t = -1.15, p = 0.26; D3: t = - 2.71, p < 0.05; D5: t = -2.62, p < 0.05; D10: t = -5.33, p < 0.001). Thus, beyond a threshold of practice, i.e. by day 3, there was a significant drop in performance in the probe trials that was not fully recovered throughout the entire probe period. Notably, a similar transient decrease in performance was also apparent on the return from the probe track to the training track, suggesting that it was not specific to the training track. The sustained decrease in performance, however, suggested a genuine limitation of participants’ ability to perform the task on the mirror-reversed track.

We suspected that the chance of falling off was not uniform along the length of the track, and so derived an alternative and more fine-grained measure of task performance using the hazard rate. The hazard rate describes the chances of falling off at each length along the track, accounting for the fact that the ant-car has already reached that length (Simes and Zelen, 1985). We found that the hazard rate was non-uniform along the length of the track (Figure 4a). We also considered the closely related survival function – the probability for a given distance along the track that participants will make it at least that far. (Survival is, essentially, the integral of the hazard rate; see Methods). Survival (Figure 4b) significantly improved across days of practice (log-likelihood ratio test between survival on days 1 & 10: X^2^(1) = 355.1, p < 0.001).

**Figure 4:**
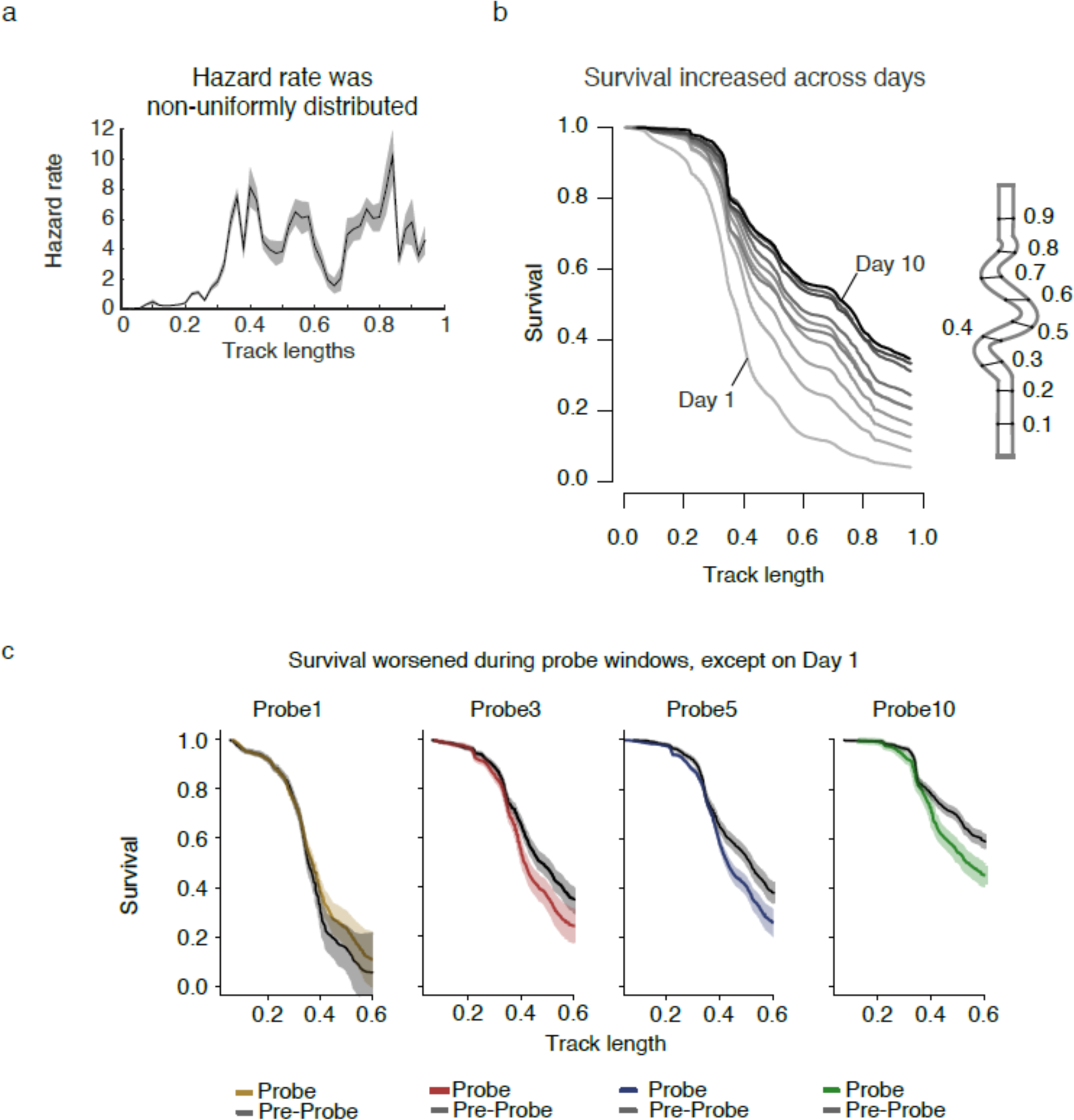
Falloff risk decreased with practice. a, The hazard rate (fall offs per 0.02 of track length) as a function of track length, pooling across all groups and days. b, The survival functions across days of training. Darker curves signify later days. c, Survival curves during blocks of Probe trials compared to Training trials on the same day as the Probe.

The survival-based analysis of performance in probe trials resulted in a similar pattern of results as the analysis based on distance travelled (Figure 4c). Survival on probe days was significantly different across the groups (log-likelihood ratio test: X^2^ = 5320.3, p < 0.001), indicating a practice benefit. Survival also differed during the probe windows compared to the pre-probe windows (log-likelihood ratio test: X^2^ = 285.42, p < 0.001), demonstrating the drop in performance during probes. There was also a significant interaction between group and probe window (log-likelihood ratio test: X^2^ =1117.1, p < 0.001).

This analysis confirmed that performance deteriorated during probes, and that the extent of this change differed significantly depending on the day it was experienced. Post-hoc tests revealed that there was no detectable difference in survival between pre-probe and probe windows for group D1 (log-likelihood ratio test: X^2^ = 0.33, p < 0.57), but there was a significant change in survival during the probes for each other group (log-likelihood ratio tests: D3: X^2^ = 50.6, p < 0.001, D5: X^2^ = 80.0, p < 0.001, D10: X^2^ = 123.8, p < 0.001). These findings are consistent with those from the distance travelled measure and confirm that task success decreased in the probe from day 3 onward.

While performance during probes experienced a consistent drop relative to pre-probe trials from day 3 onward, performance during probes nevertheless increased across days of training (linear regression: F(1, 67) = 25.9, p < .001). The fact that performance during probes improved with practice on the training track demonstrates that this continuous motor skill is generalizable. Furthermore, the fact that the performance decrement during probes was of similar magnitude after 3, 5, and 10 days of training shows that there was not an increasing divergence between the training and probe tracks that would suggest increasing task specificity. These findings are inconsistent with the theory that continuous sequential actions are generated by executing a chunked sequence of actions. Nevertheless, the fact that performance during probes did drop consistently relative to same-day performance on the training track could indicate some component of performance may have been attributable to chunking.

Purely analyzing metrics of task success based on whether and when participants fell off the track is limited since there are multiple possible reasons that a participant may have failed on a given trial, including poor selection of actions, noisy execution of actions or, alternatively, momentary lapses of control (Wichmann and Hill, 2001; Pisupati et al., 2021; Ashwood et al., 2022). Therefore, to better understand the reasons behind participants’ improved performance across days and the drop in performance on the probe trials, we analyzed the kinematics of the ant-car trajectories.

We first measured ant-car trajectory variability (Figure 5a), which we quantified as the dispersion across trials in windows of 25-trials using the first five principal components of kinematic data (see Methods). The dispersion systematically decreased with practice on the training track (Figure 5b; Linear regression; F(1, 2657) = 206, p < 0.0001). We then compared trajectory kinematics on the probe track to those on the training track. Given that performance deteriorated at the level of task success, it might be expected that kinematics would likewise revert to a level seen with fewer days of training.

**Figure 5:**
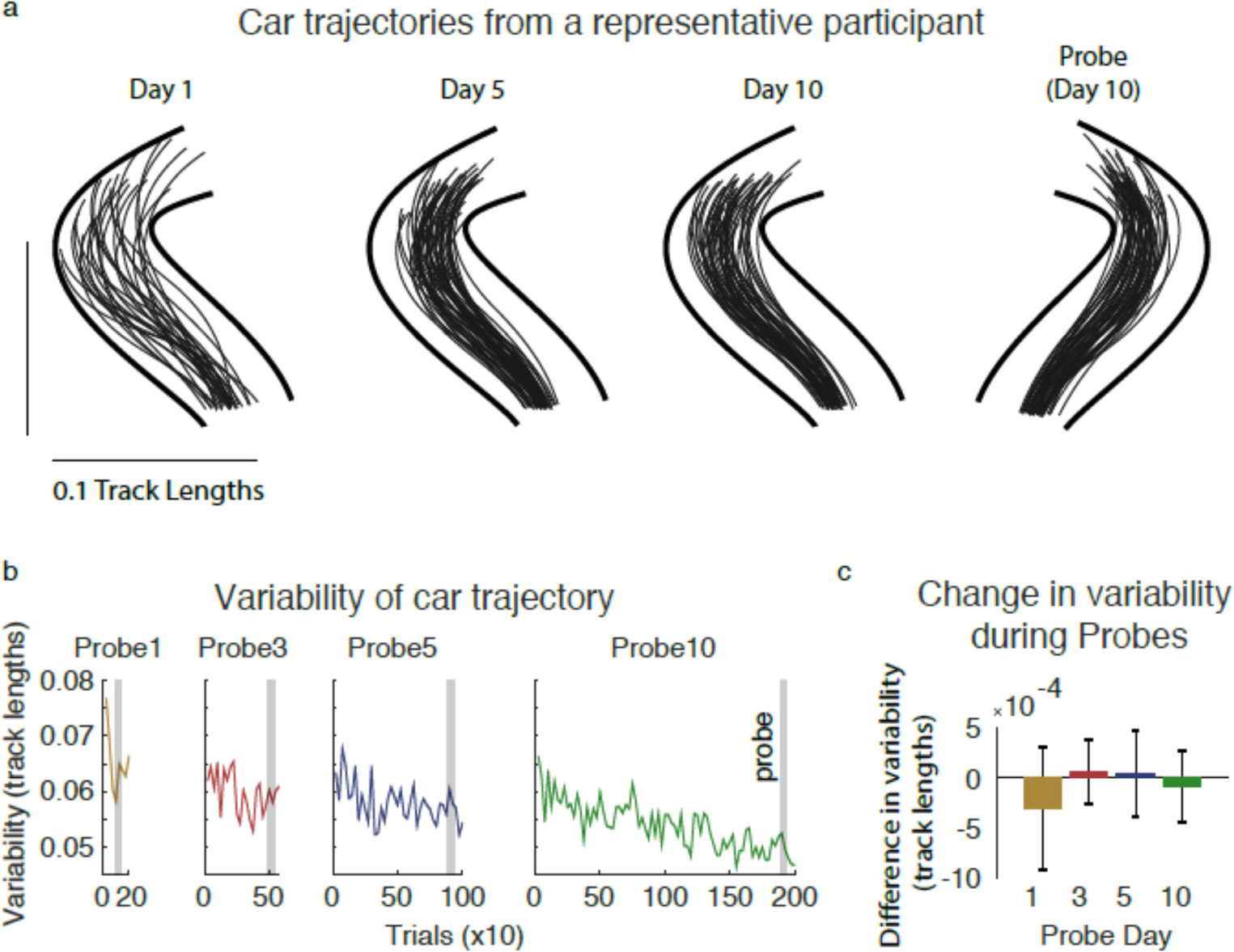
Trajectories became more stereotyped with practice. a, Sample trajectories from a portion of the Training track from a representative participant on Days 1, 5, and 10, and from a portion of the Probe track. b, Trajectory dispersion for each group across trials of practice. c, The difference in trajectory dispersion, as defined in b.

This is not what we found, however; there was no detectable difference in dispersion between probes and the pre-probe training period (Figure 5c). An analysis of variance test conducted on the difference in dispersion between the probe and pre-probe windows across groups failed to detect a difference (ANOVA: F(3) = 0.126, p = 0.94). Nor did any group individually experience a significant change in dispersion during the probe (Linear regression; D1: t = -0.600, p = 0.55; D3: t = 0.537, p = 0.59; D5: t = 0.546, p = 0.59; D10: t = 0.344, p = 0.73). However, the lack of a significant difference is not sufficient evidence for the absence of an effect of the probe. We therefore also computed the Bayes Factor (BF) of a linear model fit to the difference between probes and pre-probe windows as a function of the day at which the probe occurred, using a uniform prior (Wagenmakers, 2007). This analysis revealed a BF of 0.002, which is considered very weak evidence for there being a relationship. Note that measures of dispersion were calculated based only on successful trials. While this might at first appear circular, we emphasize that it was still possible to observe clear differences in dispersion across days using this approach and therefore, the similarity of the dispersion between probe and pre-probe windows was not inevitable. Thus, kinematic variability on the probe track did not appear to differ systematically from the training track, even though variability changed significantly across learning. Therefore, although the improvement in task success over days was consistent with a reduction in kinematic variability with practice, the drop in task success in probe trials was not attributable to increased kinematic variability.

To address the apparent paradox of there being decreased overall success on the probe track (falling off more often) even though skill at the level of kinematics fully generalized from the training track to the probe track, we more closely examined the failures. Importantly, failure trials were not included in the variability measure of skill, which was derived from only successful trials. It is possible that there was a categorical difference in training and probe trials in the nature of the failures rather than the successes. Analyzing failure trials is challenging since each failure is unique (unlike success trials which are all similar and can be easily analyzed collectively). To understand how and why failures differed from successful trials, we empirically determined participants’ state-dependent control policy based on their successes on the training track after extensive practice (on day 10). This allowed us to quantify the deviation of participants’ actions from this ideal policy throughout individual trials (Figure 6a, see methods for more details). We found that, in failure trials, there was a stereotyped increase in policy deviation just before the point of fall-off compared to successful trials at corresponding segments of track (figure 6b), suggesting that performance was perfectly good in failure trials up until a specific point where the failure began. Critically, this pattern of failure appeared to be almost identical for the training and probe tracks for any amount of practice (3, 5 or 10 days). That is to say, fall-off trials had the same kinematic form on either track; there was no detectable difference in policy deviation between the probe and pre-probe periods for any group on failure trials (t-test: D1: t = -0.92, p = .037; D3: t = -0.12, p = 0.90; D5: t = 0.67, p = 0.51; D10: t = 0.56, p = 0.58). Thus, it appears that the failures were largely attributable to momentary lapses. Indeed, there was a difference in the probability of a trial being successful between probe and Training trials for groups D3, D5, and D10 (Figure 6e; t = 1.05, p <0.01). We therefore conclude that overall performance differences between the training and probe tracks were due to momentary lapses in control occurring more frequently on the probe tracks, rather than an overall decrease in the general quality of control on the probe track.

**Figure 6:**
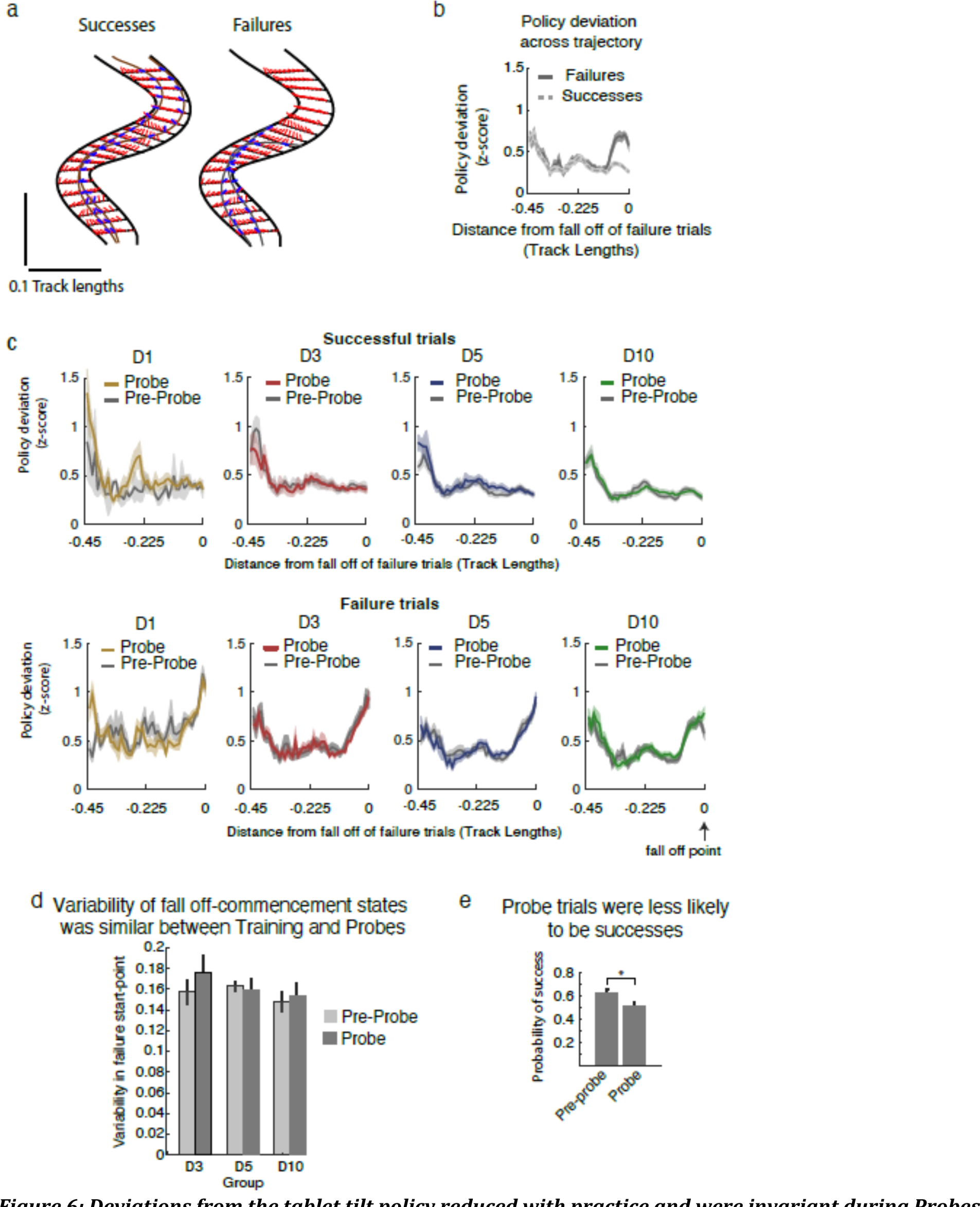
Deviations from the tablet tilt policy reduced with practice and were invariant during Probes. a, The policy was defined as the mean and variance of the tablet tilt at each discrete state among trials that successfully reached track length 0.7. The policy is shown collapsed across the multiple car direction states. Red lines indicate the average tilt direction among successful trials for each state; blue lines indicate the tilt direction for example trials, brown traces are the trajectories for those trials among successes, grey traces are the trajectories for those trials among failures. b, The deviation from policy among Successes (trials that reached at least 0.8 track lengths) and among Failures (trials that did not reach 0.8 track lengths). c, Policy deviation during Probes and Pre-Probes, broken out by day and whether the trial was successful or not. d, Variability in the location of failure start points (the track location at which the policy deviation began to diverge for failure trials). e, The probability of success for probe and pre-probe trials.

One potential objection to the idea that the increased rate of failures on the probe track was due to categorical lapses is that a small loss of skill might be apparent in a difference in the states from which failures occurred on the two tracks, i.e., they failed in a similar way but not from the same places. Thus, we tested whether the distribution of states from which failures occurred differed between the pre-probe and probe windows. We used the policy deviation signal of failure trials to identify the state from which trials began to deviate, applied principal component analysis to that distribution of multidimensional states (position and velocity of the ant-car), and computed the standard deviation of the first principal component (Figure 6d). We only applied this analysis to groups D3, D5, and D10, where we observed a sustained drop in performance on the probe track. The variance differed between the pre-probe and probe windows for group D3 only (permutation test, p < .001); for groups D5 and D10 there was no detectable difference in variance (permutation test, D5: p = 0.073, D10: p = 0.67). This suggests that the apparently abrupt onset of failures was not due to participants having gradually lost control and drifted into undesirable states; failures began at states that were also typically occupied during successful trials.

In summary, we found that there was a practice-related increase in skill across days and this skill fully generalized to a mirror track in terms of trajectory variability, control policy, and the states visited. Full generalization was masked by an overall performance asymmetry explained by the fact that subjects showed more lapses in policy compliance on the probe track than on the training track. There was no evidence for decreased generalization at the level of kinematics as skill increased, arguing against convergence on a chunked sequential action.

## Methods

### Participants

81 human participants (47 female) completed this study. All participants were 18 to 40 years of age, had no known neurological disorders, were self-reported right (76) or left (5) hand dominant, and provided informed consent to participate. The Johns Hopkins University School of Medicine Institutional Review Board approved this study and all of its procedures.

### Experimental Procedure

The study was conducted using a custom-built video game (“the game”), developed by Max and Haley, Inc. (Baltimore, MD) for the Kata Project at The Johns Hopkins University. The game simulated a driving scenario. Participants steered a virtual arthropod (“the car”) along a narrow track by tilting (i.e. changing the pitch and roll) an iPad (Apple, Inc., Cupertino, CA) computer (Figure 1a). The direction of the acceleration of the car was obtained by projecting the vertical axis of a world-centered coordinate system onto the tablet’s surface, giving a magnitude and direction vector; which, by analogy, would be the direction and magnitude of acceleration of a marble rolling off of a flat surface if tilted. The kinematics of the car in the game were obtained from a physics simulation that included the interaction of the multiple car segments, which introduced nonlinearity in the mapping between the tablet tilt input and the car’s dynamics. These computations acted as a filter that introduced a delay of approximately 50 ms between the tablet tilt and the response of the car. The magnitude of the tablet tilt vector was set to a constant value, making the tilt magnitude a control null-space and effectively constraining the speed of the car in the game to a narrow range. The game’s software had a frame rate of 60 Hz, and recorded the magnitude and direction of the tablet tilt and the path of the car along the track (Figure 1b) at 60 Hz.

The experiment included two tracks: a training track and a probe track (Figure 1c). The probe track was the mirror image of the training track. This guaranteed that the two tracks were matched for difficulty and that successfully navigating each track would require unique actions in a novel sequence relative to one another (Supplemental Figure 1). The track that was designated as probe or training was counter-balanced across participants in each group.

Participants were assigned to one of four possible groups. Groups differed in the number of days of training that were conducted using the training track before the probe track was introduced (Figure 1d). Groups D1, D3, D5, and D10 experienced the probe track on the first, third, fifth, or tenth day of training, respectively. Groups D3 and D5 each conducted five total days of training. Group D1 completed the study after the first day of participation, and group D10 completed the study after 10 days of participation.

Each day of practice included 200 trials and lasted approximately 30 minutes. Trials in which the entire track was completed lasted approximately 5s. A 4s delay was imposed if the car fell from the track, which would happen if the track’s edge was breached. Inter trial intervals (the time between the successful completion of one trial and the beginning of the next, or the time after the 4s-delay of a failed trial and the beginning of the next trial) lasted 3s on average and were self-paced; participants pressed a button on the device’s screen to begin the next trial. The car’s dynamics were invariant for the duration of the experiment including on the probe Track. Probes consisted of a block of 50 contiguous trials in which the probe track was attempted instead of the training track. Participants were not pre-warned that a probe block would be experienced. 67 participants took part in the study in the BLAM laboratory at the Johns Hopkins Hospital, and 14 had the game downloaded onto their personal iPad devices and completed training for the study from home. All sessions that included the probe track were conducted in the laboratory using the same individual iPad on which each participant trained.

### Data Analysis

Data were analyzed offline using Matlab (The Mathworks, Natick, MA, 2013) and R (The R Project, www.r-project.org). All code is available online at https://github.com/dhuberdeau/iPadGame. For each trial, the position along the track at which the car fell off was detected by searching for breaches of the track boundary. The length of track that the car reached by the fall off point was recorded in units of the fraction of the total track length, a quantity between 0 and 1.

The fall-off hazard was assessed as a function of the length of the track. The hazard rate of car falloffs, ²A, as a function of track distance, *t*, is given by the conditional probability

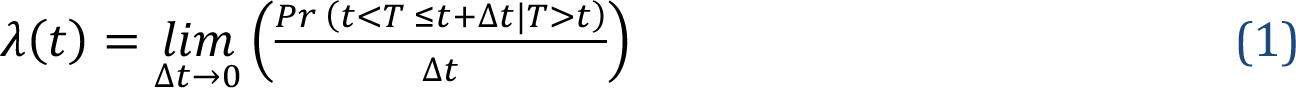

where *T* is a continuous random variable representing the track length at which a car fall-off event occurred. Suppose that *T* has the *pdf*, or probability density function, 𝑓(𝑡), and *cdf*, or cumulative distribution function, *F*(*t*), then the hazard rate function is related to the *pdf* and the survival function, 𝑆(𝑡) = 1 − 𝐹(𝑡), by the following equation.

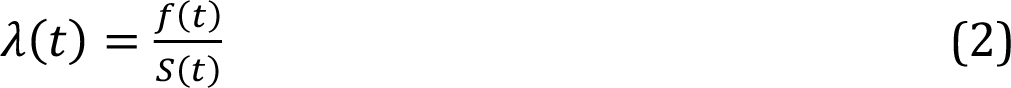

An estimate of the survival function for each participant on each day of Training and during probes was obtained using the Kaplan-Meir method (Borgan, 2001).

To analyze movement kinematics, a measure of the variability among car paths across trials was computed for each participant. A segment of each trial’s car path was isolated from the time at which the car reached track length 0.25 and for 750 ms thereafter. Only trials for which a fall off did not occur prior to or during this window of time were included in the analysis of kinematics; we refer to such trials as qualifying. In order to compare kinematics across participants that were assigned different track orientations, and to compare between training and probe conditions, any car paths that used the orientation depicted in Figure 1 as the probe track were flipped across the vertical axis to match the training track.

The across-trial kinematic variability was computed as the dispersion (see Equation 3) of the first five principal components after applying principal component analysis decomposition to the car’s path. For each participant, all qualifying trajectories from all other participants were pooled together (a leave-one-out approach at the level of participants) in order to form a basis. Trajectories from the given participant were then projected into this basis. The top five principal components reliably accounted for over 99% of the variance in the data for each participant. All qualifying trajectories from each participant were projected onto the axes corresponding to the first five principal components, and the dispersion *d* of these samples was computed by taking the sum of the Euclidean distance between each pair of distinct samples (x_i_ and x_j_) and dividing by the number of pairs.

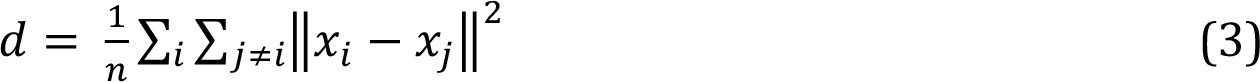

Any window of trials that had fewer than seven qualifying trajectories was excluded from further analysis.

Another analysis was developed to measure the extent to which the tablet tilt signal on a given trial deviated from the optimal policy. The track was discretized along its length into 90 bins, across its width into 30 bins, and across car direction headings into 100 bins. An empirical state-dependent policy was computed that consisted of the average tablet tilt direction at each state from among those trajectories that were ultimately successful (Figure 6a). Trials were labelled successful if they reached at least to track distance 0.8. The policy map was generated for the region of track between lengths 0.25 and 0.75, a region that included the first two turns in the track and the first two peaks of the hazard rate. A policy map for each individual was computed by pooling the kinematic data from Day 10 of participants in group D10, including using a leave-one-out approach for participants in that group. Only D10 participants were used to form the policy because this group experienced the most training of all groups and were therefore assumed to have behavior closest to a theoretical optimum. The policy deviation was then taken to be the difference between the tablet tilt direction and the empirical policy at each state visited in a trajectory.

A comparison of behavior during probe and training conditions was done with respect to distance travelled, hazard rate, car path kinematics, and policy. These comparisons were done by testing for changes in each measure between the probe and pre-probe windows of trials, excluding the first 5 trials of the probe. The pre-probe window included the 50 trials immediately before the probe, and the probe window included trials 6 to 50 of the probe, which itself lasted for 50 contiguous trials.

### Statistical analysis

The distance travelled, kinematic variability, and policy deviation, were each used to test for practice-related changes in behavior, and for differences in behavior between training and probe tracks. Practice-related changes in behavior were assessed independently for each metric by fitting a linear model to the average of that metric within windows of 25 trials. Differences between training and probe trials were assessed first by conducting an analysis of variance test on the within-subject difference in mean of each metric between the probe window and the pre-probe window, with group as the independent factor. In the event of a significant group difference, independent pair-wise t-tests were conducted to determine which group(s) differed from one another. An additional analysis was conducted for each group to determine whether the difference in metric between the probe and pre-probe windows was significantly different from zero. To test for changes in survival between the probe and the pre-probe window, a proportional hazard model was fit to data from each window and tested for changes using a cox mixed-effects model. All statistical analyses were conducted in R (www.r-project.org).

## Discussion

Here, we created a novel video game to investigate how a movement skill made up of continuous sequential actions is acquired through practice. We reasoned that if participants chunk a series of discrete actions (such as the tablet tilts needed to successfully complete the turns of the track) then they would show poor generalization to a mirror-image version of the track that required different actions to navigate. If, instead, they learned a flexible control policy, then they would generalize. We found that practice over 10 days led to improved task success and a reduction in trajectory variability when steering along a curved track. There was full generalization to the mirror-image track at the level of kinematics but not task performance. The performance difference found between the training and probe tracks after day 1 was not due to a failure of skill generalization but to a greater likelihood to lapse from one’s skilled state on the less familiar probe track. We conclude that continuous sequential actions are learned as control policies that map states onto actions.

In previous work, we have shown that for a task that required fast continuous semi-circular movements of the wrist (arc pointing task, APT), participants got better over many days of practice (Shmuelof et al., 2012). This improvement was measured at the level of task success as a shift in the speed-accuracy trade-off and at the level of execution as smoother and less variable movement trajectories. A critical question is what kind of representation supports practice-driven improvements in kinematic performance of the kind observed in the APT. In the sequence-learning literature, the vast majority of which has been about discrete tasks, a prominent idea has been that of chunking, whereby each individual movement gets incorporated into single larger motor unit (a chunk) that can then be expressed all at once (Ramkumar et al., 2016; Krakauer et al., 2019; Yokoi and Diedrichsen, 2019; Berlot et al., 2020). It seems intuitive when looking at the evolution of the continuous wrist movements in the APT that they too went from a series of faltering sub-movements to a rapid single swipe. Findings in that study, however, suggested otherwise. First, kinematic analysis revealed that although the trajectories through the tube became smoother and less variable with practice, sub-movement number remained invariant. Thus, the skill comprised better concatenation of the execution primitives rather than fusing them into a single movement. Second, we found evidence for improved feedback corrections; in the case when the trajectories got too close to the edge, more practiced subjects showed superior ability to steer away. These two results from our former study were clues that perhaps subjects were not just chunking sub-movements into a stereotyped trajectory but instead were learning a more effective feedback control policy, which would also reduce trajectory variability and thus give the appearance of stereotypy. These conclusions were provisional, however, and a follow-up study was required to support them.

Here, in the ant-car game we saw a similar reduction in trajectory variability along the track as was seen in the APT. Additionally, however, we were able to show full generalization to the probe track at the level of kinematics, which rules-out chunking and is consistent instead with acquisition of a flexible feed-back control policy. Generalization was not due simply to a concordance between the two tracks. An analysis of the relative similarity of the two tracks revealed that they were dissimilar, and thus specific sequences or subsequences of actions that would lead to success in one were unlikely to be successful in the other if applied verbatim.

Based on the results here, we suggest that increases in skill in continuous control tasks do not occur through selecting and combining movements into a sequence of actions that can then be subsequently chunked (Johnson, 1970; Robertson, 2007; Wong et al., 2015; Wong and Krakauer, 2019; Yokoi and Diedrichsen, 2019). Instead, a novel feedback control policy must be learned from scratch and applied to a continuous sequence of states. The learning of such *de novo* control policies is distinct from adaptation and discrete sequence learning because it requires both rapid selection of a new response and proper execution of that response (Yang et al., 2021). Thus, we would conjecture that chunking in motor learning only occurs in those tasks that allow for an overt abstract or cognitive representation at the level of action selection. For example, one can rapidly press the keys of a bank cash machine because overt knowledge of the passcode has been chunked. This cognitive chunk is then fed to a motor area for rapid serial execution of over-learned finger movements (Wong and Krakauer, 2019; Yokoi and Diedrichsen, 2019; Zimnik and Churchland, 2021). Chunking is not an available strategy for motor tasks that are not conducive to a cognitive action representation phase and when motor execution itself must improve.

An interesting aspect of our study was that participants consistently showed deterioration in performance at the level of task success on the probe track – a 10% drop from day 3 onwards. There was a need to resolve the apparent contradiction between full generalization at the level of execution but not at the level of overall task performance. To do this, we derived an empirical control policy from the best mean performance on the training track at day 10. The usefulness of the measure is that it is independent of which specific states are visited on the track, which may be suboptimal, but in any given state there is nevertheless an optimal policy to follow. We reasoned that full generalization would mean that participants could transfer this policy to the probe track: given a state (position and velocity) on the track there is an optimal tilt angle. Interestingly, this is indeed what we found; the policy profiles for successful trials were the same on both tracks. One remaining difference, however, might have been that although the same policy was being used at any given position on either track, participants might have been choosing a different trajectory on the probe track compared to the training track, i.e., not the mirror image. To address this we looked at the distribution of states visited on the two tracks, and they were not different. The full transfer of the learned control policy from the training track to the probe track corroborated the finding of an equivalent degree of practice-related variability reduction on both tracks. Overall, there was no evidence for any qualitative difference in skill when participants steered the car on the probe versus the training track.

The policy deviation metric also allowed us to search for an alternative explanation for why there were more failures on the probe track given that participants had the same level of skill as on the training track. For the training track, visual inspection of the unfolding of the control policy in trials where a fall-off occurred compared to neighboring trials that did not fail, revealed that failures occurred through an abrupt error in control immediately prior to the fall-off; control looked like successful trials right up to that point. Critically, just like in the case of successful trials, the failure policy profiles on the probe track were the same as those on the training track. The difference in overall performance between the two tracks was instead attributable to an increase in the probability of making an abrupt error leading to a fall-off, i.e,. a *lapse*. Thus the probe track did not induce failure to generalize skill but instead caused an increase in the probability of lapses of the same kind that occurred on the training track.

Chomsky famously made the performance versus competence distinction in the context of language (Chomsky, 1965). Performance can be hampered by false starts and slips of the tongue, but this does not indicate loss of knowledge of the language. We conjecture that we saw the analogous situation here: participants had equal skill (competence) on each track, but exhibited more lapses, measured as a performance drop in terms of task success. Lapses are ubiquitous in behavioral data and describe instances in which mistakes are made even when they are predicted not to occur, such as in perceptual decision making tasks in the presence of strong evidence for one decision outcome over another (Wichmann and Hill, 2001). Lapses have been hypothesized to arise from inattention or from errors of execution, such as motor noise. However, a more recent suggestion is that they indicate targeted exploration in the presence of uncertainty (Pisupati et al., 2021) or deliberate alternation between behavioral strategies (Ashwood et al., 2022). While lapses have been rarely studied or discussed for motor behaviors, any of these putative mechanisms for lapses could account for the results we found here, such as inattention, strategic exploration in the presence of a novel set of states, or even possibly a reduction in motivation (Wong et al., 2015).

Here we draw two main conclusions. First, skilled execution of a continuous sequential action is achieved by *de novo* learning of a feedback control policy that maps states onto actions and not through chunking of a sequence of actions. This suggests a fundamental difference in how motor skills are acquired when they require improvements in movement execution versus those that require selection between movement elements that can already be well executed. Second, performance can vary across contexts for the same level of skill because of differences in the frequency of lapses. This is intriguing and potentially profound, as it suggests that motor skills are only as good as our ability to consistently express them. This is well known in the sports world; there are athletes recognized for their brilliant skill, but they seem unable to maintain their highest-level performance from one game to another.

## Acknowledgements

We thank Dr. Omar Ahmad and Promit Roy for programming the antcar game. This work was supported by funds from The Johns Hopkins University Science of Learning Institute.

## Supplemental Materials

We sought to quantify the extent to which tablet tilt signals for successful trials were similar between the probe and training tracks. Highly similar tablet tilt signals could potentially account for any generalization of performance across track types, while dissimilar signals would make generalization on account of the similarity between the tracks themselves less likely. Signal similarity was computed as the Euclidean distance between a segment of the tablet tilt signal from each probe trial and the best matched segment of equal length from among tablet tilt signals of training trials. The window lengths used to isolate signal segments were 0.1s, 0.2s, 0.3s, 0.5s, 1s, and 2s. Signal similarity was computed across the length of the track in step sizes of 33% of the width of the window. Only trials that had completed the track were included in this analysis. Since segment window size affects the Euclidean distance, relative signal difference was computed as the ratio of the Euclidean distance between the training and probe signals to the Euclidean distance between training trials taken before and after the probe (inverting the pre-probe signal so that it matched the signals from probe trials). Large Euclidean distances between probe and training tracks would signify that the two track types were not similar, and thus would rule out the possibility that any generalization of performance during probe trials was due to similarity between the tracks.

**Supplemental Figure 1:**
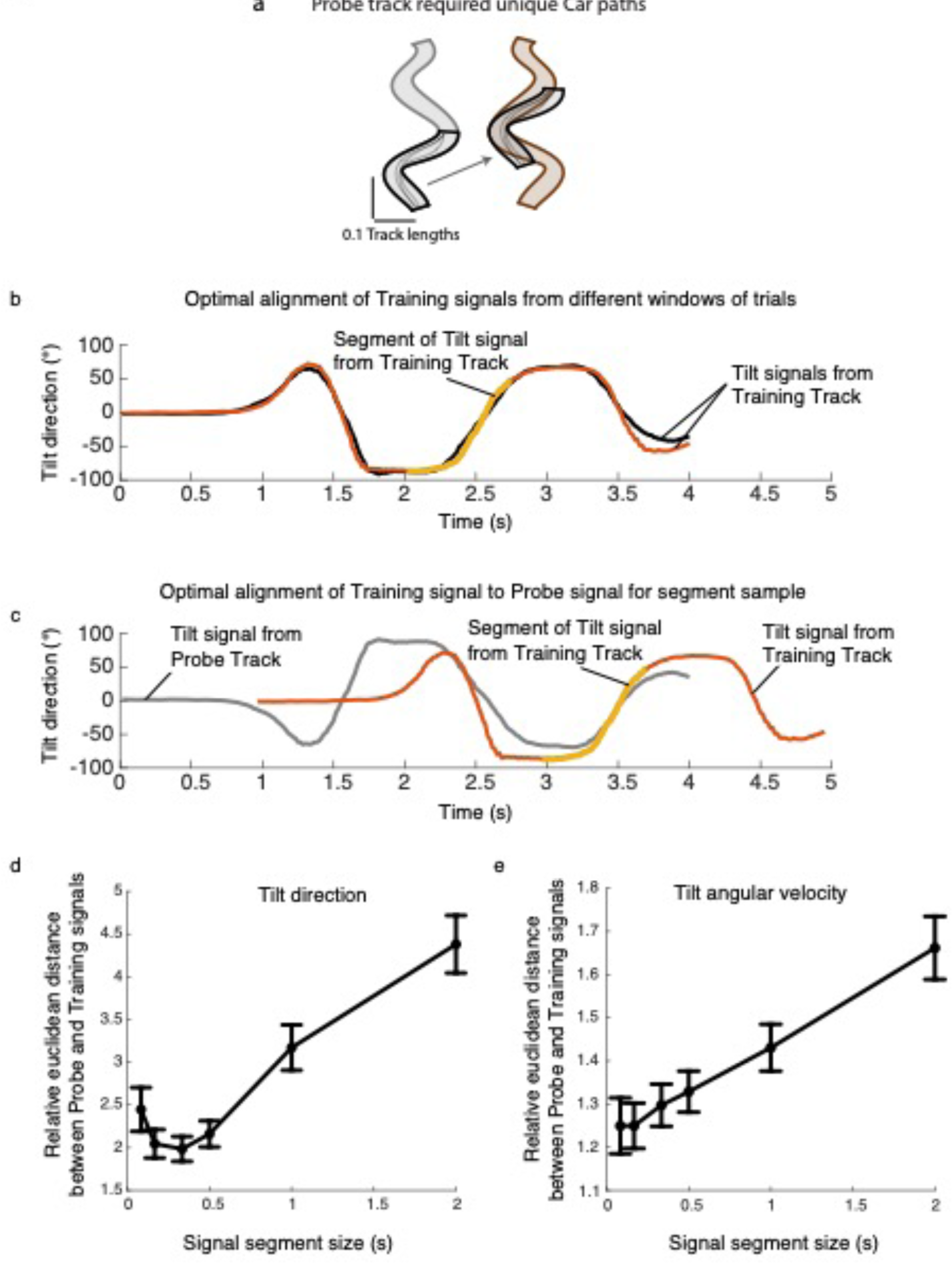
Probe track required different kinematics than the Training track. a, The two tracks used as Probe or Training. The analysis of kinematic similarity attempts to determine if segments of kinematics that would be successful in one track would also be successful in the other. b, Optimal match of a segment of kinematics (yellow highlight) from a tablet tilt signal from the Training track against the average Tablet tilt signal of Training tracks. c, Optimal match of a segment of kinematics (yellow highlight) from a tablet tilt signal from the Probe track against the average Tablet tilt signal of Training tracks. d, The normalized euclidean distance between Probe track kinematics and Training track kinematics of the tilt direction signal. e, As in d but for the derivative of the tilt direction signal

